# Regulating of Koisio technology-produced fluid on Vascular Adventitial Interstitial Fluid flow and its Effect on heart rate by delivering Esmolol

**DOI:** 10.1101/2024.11.11.622840

**Authors:** Bei Li, Jin Cai, Chao Ma, Jun Hu, Hongyi Li

## Abstract

Acupoint-originated interstitial fluid (ISF) circulatory network along the vasculature has been demonstrated to play a role as an extravascular drug delivery pathway. We aim to reveal the regulation of Koisio technology-produced fluid (KF), which is rich in Ultra-small Particle Size Nanobubbles (UsNBs), on vascular adventitial ISF flow and its Effect on heart rate by delivering Esmolol. In animal experiments, using real-time fluorescent imaging technology, it was found that: when normal saline-fluorescent liquid was given at Kunlun acupoint, the velocity of the adventitial ISF along the femoral vein was measured to be about 600 μm/sec, and the width of the tunica ISF channel on the adventitia surface of the femoral vein was about 210μm; when KF-fluorescent liquid was injected into the subcutaneous tissue of Kunlun acupoint, the velocity of the adventitia ISF slowed down by about 24% compared with before; the width of the tunica ISF channel on the adventitia surface of the femoral vein increased by about 35% compared with before; When 150μL of KF with high-concentration UsNBs was injected into Kunlun acupoint, it was observed that the adventitial ISF flow velocity was decreased by 43%, and recovered after 30 minutes. Administration of normal saline-esmolol solution at Kunlun acupoint caused the heart rate to drop from 410 bpm to 380 bpm, a decrease of 7.3%, which continued for 5 minutes before recovery. Administration of KF-esmolol solution at Kunlun acupoint caused the heart rate to drop from 410 bpm to 371 bpm, a decrease of 9.5%, which continued for 15 minutes before recovery. Thus, acupoint injection of KF can regulate the velocity of the adventitia ISF flow along the femoral vein and increase the width of the tunica ISF channel. These results may reveal a new mechanism of the physiological effect of Koisio drinking water rich in ultra-small particle size nanobubbles. KF may not only regulate the acupoint-originated ISF circulatory system, but also play a role in adventitial drug delivery, nutrient supply, and waste removal.

## Introduction

Acupoint-originated interstitial fluid (ISF) circulatory network along the vasculature has been demonstrated to play a role as an extravascular drug delivery pathway, and potentially enhance therapeutic effects while minimizing side effects ^1^. Previous study has shown that the acupoint-originated ISF circulatory network is part of the ISF circulatory system which is distributed all over the whole body ^1^, with the characteristics of “connecting acupoints in the extremities, in the adventitia along the vasculature, and targeting the heart for ISF flow and perfusion system”. It has the potential to become an extravascular drug delivery pathway and has drug delivery and metabolism characteristics different from intravenous blood circulation ^2^. It is expected to achieve targeted drug delivery to organs such as the heart and lungs, thereby reducing drug dosage and drug side effects potentially. Animal experiments have confirmed that substances such as SDS that can change the surface tension of liquids can significantly change the flow of vascular adventitial ISF^3^, and are expected to become a way to regulate drug delivery and efficacy. In the past three decades, nanobubbles have not only been found to be able to exist stably for hours or even days, but also have special properties such as high density^4^, large specific surface area^5^, and changing surface tension^6^. Nanobubbles have gradually been widely used in various fields such as agriculture, industry, and biomedicine. In this context, the use of Koisio technology-produced fluid (KF) which is rich in Ultra-small Particle Size Nanobubbles (UsNBs), provides a promising way to regulate the drug delivery of the acupoint-originated ISF circulatory system. This study explored the regulation of KF on vascular ISF flow and its Effect on heart rate by delivering Esmolol.

### Materials and Methods Animals

All animal experiments were approved by the Institutional Animal Care and Use Committee of the Institute of Basic Medical Sciences Chinese Academy of Medical Sciences, Peking Union Medical College (No. ACUC-A02-2021-017) and performed according to institutional guidelines and research protocols approved by the Institutional Animal Care and Use Committee of the Institute of Basic Medical Sciences Chinese Academy of Medical Sciences, Peking Union Medical College. A total of 27 male and 27 female Sprague—Dawley rats, 20-25 weeks old, 300-350g in weight (HFK Bio-Technology, Beijing, China), were used and housed in pathogen-free conditions with 12 hours of continuous light and 12 hours of continuous darkness at the Laboratory Animal Center of the Institute of Basic Medical Sciences Chinese Academy of Medical Sciences. Equal numbers of male and female rats were taken to ensure that sex of the animals does not constitute a biological variable during analysis. The procedures designed minimize animal suffering and respect the 3Rs principles. The rats were anaesthetized with isoflurane (1%) in 1 L/min oxygen or intraperitoneally anaesthetized (pentobarbital 50 mg/kg) during experiments. The body temperature was maintained at 37.5 °C with a rectal probe-controlled heated platform. The rats were euthanized by carbon dioxide according to the guidelines of the American Veterinary Medical Association (AVMA).

### Kunlun acupoint injection of KF

A total of 54 rats were randomly divided into 9 groups, with 6 rats in each group. The rats were injected with corresponding concentrations and volumes of UsNBs or Ultra Purified Water (UPW) at Kunlun acupoint.

Preparation of KF was described according to previous work^7^. The KF with low concentration UsNBs (LC-UsNBs, 10^7^-10^8^ particles/mL) or high concentration UsNBs (HC-UsNBs, 10^10^-10^11^ particles/mL) was produced by the technical experts of Shanghai Koisio Food Industry Co. (Shanghai, China) according to standard procedures.

### Real-time fluorescence imaging of adventitial ISF flow along femoral vein

After the skin along the vessels of the limbs opened surgically, the Fluorescein Sodium (FluoNa) was administrated by adventitial infusion on the saphenous vessels into the saphenous vein. The real-time continuous speckle-like flow of the fluorescent adventitial ISF along the arterial and venous vessels in the limbs was dynamically recorded by the fluorescence stereomicroscope (Zoom V16, Carl Zeiss, German) with a high-sensitivity camera (Prime BSI Scientific CMOS, Teledyne, USA).

### Measuring fluorescent ISF flow velocity and width

The fluorescent ISF flow velocity was measured by a method of Speckle tracking Velocimetry (STV) described in our previous study^2^, which has been extensively used in measuring displacements and velocities.

The fluorescent ISF flow width was calculated by using the image annotation and statistical calculation functions of ZEN 3.2 software.

### Adventitial infusion of Esmolol and heart rate recording

Esmolol was dissolved in KF with LC-UsNBs, KF with HC-UsNBs, or normal saline (NS) respectively. Consistent with previous studies^2^, The dosage of Esmolol for adventitial infusion was 0.5 mg/kg and adjusted according to each rat weight. Real-time electrocardiogram (ECG) and heart rate were recorded using animal physiological monitoring device (PowerLab, AD Instruments, Australia).

## Results

### KF slowed down adventitial ISF flow velocity along femoral vein

Study on the regulatory effect of KF on the acupoint-originated ISF circulatory system by injecting KF into the Kunlun acupoint and observing the changes in the adventitial ISF flow velocity along femoral vein. Injection of KF with HC-UsNBs, KF with LC-UsNBs or NS induced different changes in the adventitial ISF flow velocity. There was no change in the adventitial ISF flow velocity along the femoral vein when 50 μL or 150 μL NS were injected, respectively. When 50 μL or 150 μL of KF with HC-UsNBs or LC-UsNBs were injected, the adventitial ISF flow velocity dropped to the lowest at 5 min and then gradually recovered, with differences in the decline and recovery time. Regardless of the concentration, the larger the volume injected, the greater the decrease in flow velocity and the longer the recovery time. When 150 μL KF with HC-UsNBs were injected, the adventitial ISF flow velocity decreased the most (43%) and recovered the longest (recovered to the initial level at 30 min after injection) (Figure 1A, 1B).

**Figure 1.**
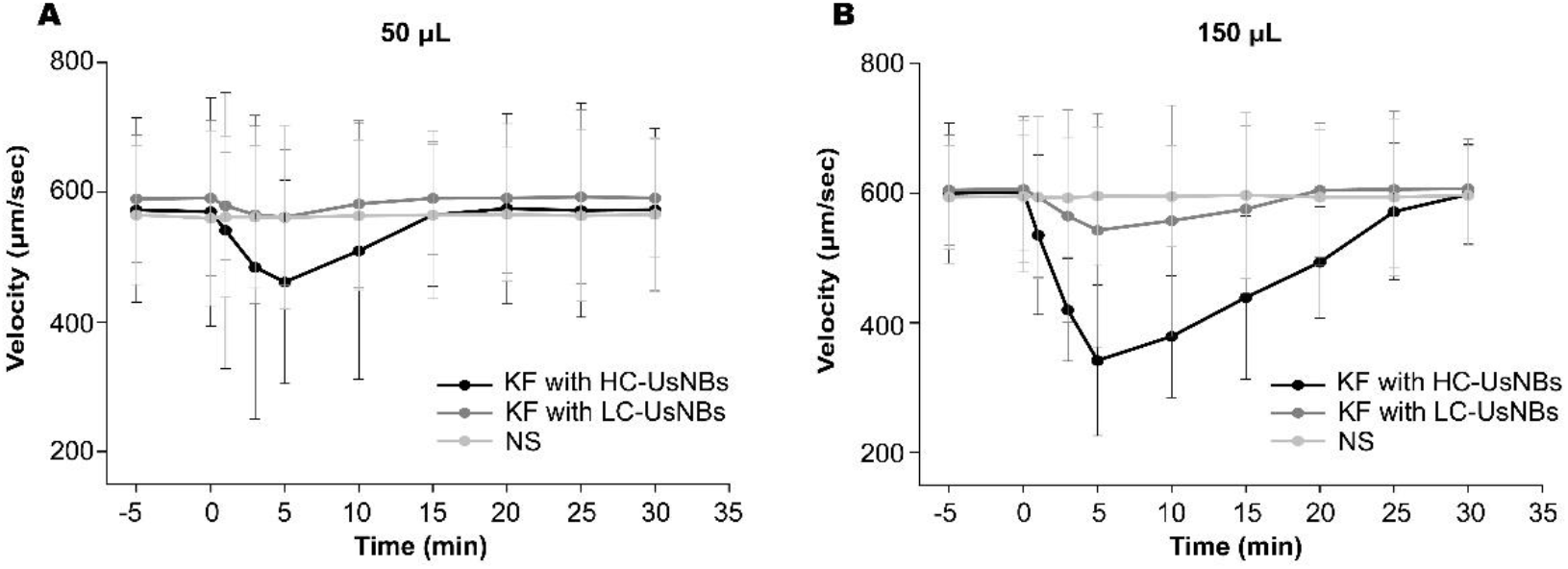
Injection of KF slowed the adventitial ISF flow velocity along the femoral vein. **(A)** Real-time adventitial ISF flow velocity along the femoral vein after injection of 50 μL KF with HC-UsNBs, KF with LC-UsNBs, or NS. **(B)** Real-time adventitial ISF flow velocity along the femoral vein after injection of 150 μL KF with HC-UsNBs, KF with LC-UsNBs, or NS. In (A) and (B), mean ± SEM, n = 6 rats.

### KF increase adventitial ISF flow volume along femoral vein

The width of the adventitial fluorescent ISF along the femoral vein after injection of KF reflected the flow volume of substances transported by the adventitia ISF. Like Results 1, injection of KF with HC-UsNBs, KF with LC-UsNBs or NS induced different changes in the adventitial fluorescent ISF flow width. There was no change in the adventitial fluorescent ISF flow width along the femoral vein when 50 μL or 150 μL NS were injected, respectively. When 50 μL or 150 μL of KF with HC-UsNBs or LC-UsNBs were injected, the adventitial fluorescent ISF flow width increased to the maximum at 5 min and then gradually recovered, with different increase amplitudes and recovery time. When 150 μL KF with HC-UsNBs were injected, the adventitial fluorescent ISF flow width increased the most (49%) and recovered the longest (recovered to the initial level at 30 min after injection) (Figure 2A, 2B).

**Figure 2.**
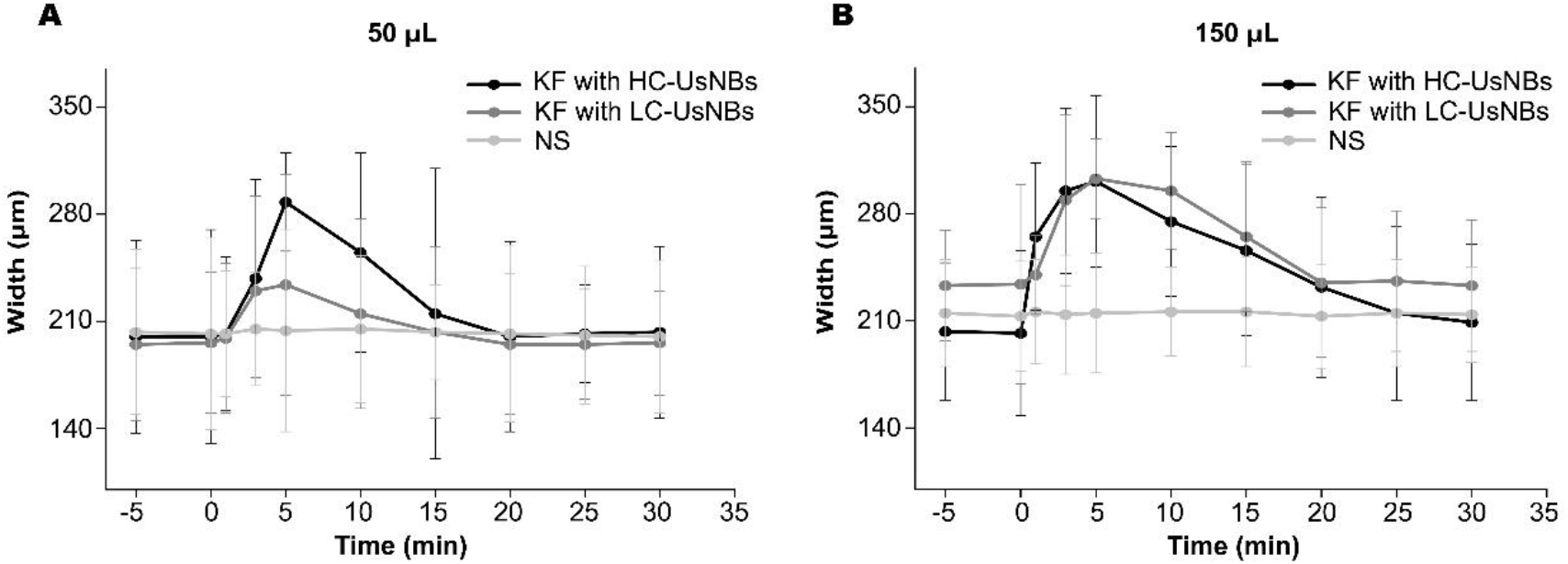
Injection of KF increased the adventitial ISF flow volume along the femoral vein. **(A)** Real-time adventitial fluorescent ISF flow width along the femoral vein after injection of 50 μL KF with HC-UsNBs, KF with LC-UsNBs, or NS. **(B)** Real-time adventitial fluorescent ISF flow width along the femoral vein after injection of 150 μL KF with HC-UsNBs, KF with LC-UsNBs, or NS. In (A) and (B), mean ± SEM, n = 6 rats.

### KF prolong the duration of esmolol efficacy

The time, amplitude and recovery time of heart rate decrease in the NS solution group were consistent with the previous study^2^. Compared with the NS solution group, in KF with HC-UsNBs or LC-UsNBs solution group, the heart rate decrease was increased and maintained for a longer time, while the time for the heart rate to return to the initial level was consistent. Moreover, the heart rate decreased more (9.5%) and lasted longer (10 min) after in KF with HC-UsNBs solution group. (Figure 3)

**Figure 3.**
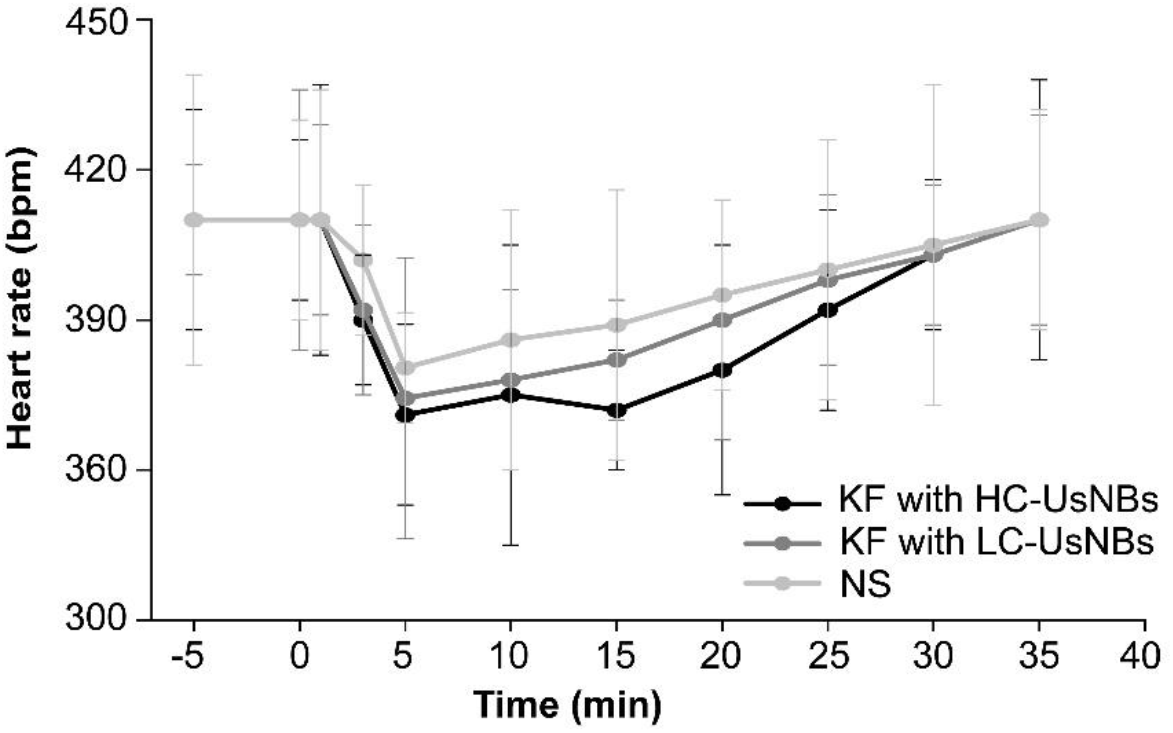
KF to dissolve Esmolol prolong the effective duration by adventitial infusion. Real-time recording of heart rate after adventitial infusion of esmolol dissolved in KF with HC-UsNBs, KF with LC-UsNBs, or NS. Mean ± SEM, n = 6 rats.

## Discussion

Previous studies have found that the acupoint-originated ISF circulatory system can serve as a unique extravascular drug delivery pathway targeting the heart and lungs, with drug metabolism characteristics different from blood delivery, and has the advantages of reduced drug dosage and long duration. Studies have found that nanobubbles can reduce the surface tension of liquid boundaries and thus affect liquid flow rate and volume, but the impact on the acupoint-originated ISF circulatory system has not been clarified. In this study, by injecting different volumes and concentrations of KF with UsNBs at the Kunlun acupoint, the changes in the adventitial ISF flow velocity and volume of the femoral vein and the characteristics of drug delivery were compared, and it was found that KF can improve the drug effect and prolong the action time by slowing down the velocity and increasing the volume of the vascular adventitial ISF flow.

### Effect on Adventitial ISF Flow Velocity

Injection of different volumes of KF with UsNBs led to changes in the adventitial ISF flow velocity along the femoral vein. KF with high concentration or low concentration UsNBs induced a significant decrease in flow velocity, with a maximum decrease observed at 5 minutes post-injection. The larger the volume of KF injected, the greater the decrease in flow velocity and the longer the recovery time. The most significant decrease (43%) in flow velocity was observed with the injection of 150 μL KF with HC-UsNBs, which recovered to the initial level after 30 minutes.

### Effect on Adventitial ISF Flow Volume

KF injection influenced the flow volume of substances transported by the adventitial ISF along the femoral vein, as indicated by changes in the adventitial fluorescent ISF flow width. Similar to the velocity results, KF with HC-UsNBs or LC-UsNBs caused an increase in the flow width, with the most significant increase (49%) seen with 150 μL KF with HC-UsNBs injection. The recovery time for the flow width to return to the initial level was 30 minutes post-injection.

### Esmolol effect on lowering heart rate

The study assessed the effect of KF with UsNBs on the heart rate-lowering effect of esmolol. Compared with NS as the solvents, using KF to dissolve Esmolol can extend the time of heart rate reduction by about 10 minutes.

In summary, our findings showed that KF with UsNBs can modulate adventitial ISF flow velocity and increase the width of the tunica channel along the femoral vein, which may reveal a new mechanism for physiological effects of UsNBs fluid. The administration of KF via acupoint-adventitia injection can reduce the flow rate of adventitial ISF and potentially expand the diameter of tunica channels along the veins. Meanwhile, the administration of KF plus Esmolol via acupoint-adventitia injection prolonged the duration of heart rate reduction, which is probably due to the decrease in the flow of adventitial ISF along the vasculature or even an increase in the diameter of the blood vessel. These findings indicate that KF might play a role in regulating acupoint-originated ISF circulatory system, and modulating the drug delivery, nutrients supply or waste products clearance along the vasculature.

## Author contributions

HY.L. conceived and developed the original ideas, concepts and theory of interstitial fluid circulatory system. HY.L and J.H. designed the current experiment. B.L. and J.C. performed the experiment. B.L. wrote the paper.

